# Magnetite Nanoparticle Photothermal Therapy in a Pancreatic Tumor-on-Chip: A Dual-Action Approach Targeting Cancer Cells and their Microenvironment

**DOI:** 10.1101/2025.02.06.636875

**Authors:** Anastasiia Dubrova, Charles Cavaniol, Aurore Van de Walle, Paul Mathieu, Zoé Fusilier, Nader Yaacoub, Yoann Lalatonne, Stéphanie Descroix, Claire Wilhelm

## Abstract

The application of magnetite nanoparticles (MagNPs) for photothermal therapy (MagNP-PTT) has recently expanded in cancer treatment. This study introduces MagNP-PTT in a tumor-on-chip model to target highly aggressive pancreatic ductal adenocarcinoma (PDAC). A tumor-on-chip system was developed using PANC-1 PDAC cells embedded in a collagen type I extracellular matrix and cultured for one week to form tumor spheroids. This platform serves as the foundation for applying PTT in a model system that is aimed to mimic the native tumor microenvironment. MagNPs efficiently penetrate the tumor spheroids, achieving controlled heating via near-infrared (NIR) light. By adjusting nanoparticle concentration and laser power, temperature increments of 2°C between 38–48°C were established. Temperatures above 44°C significantly increased cell death, while lower temperatures allowed partial recovery. Beyond inducing cancer cell death, MagNP-PTT altered the extracellular matrix, and triggered a slight epithelial-mesenchymal transition marked by increased vimentin expression. These findings highlight MagNP-PTT as a dual-action therapy, targeting both tumor cells and their microenvironment, offering a novel approach for overcoming stromal barriers in pancreatic cancer treatment.

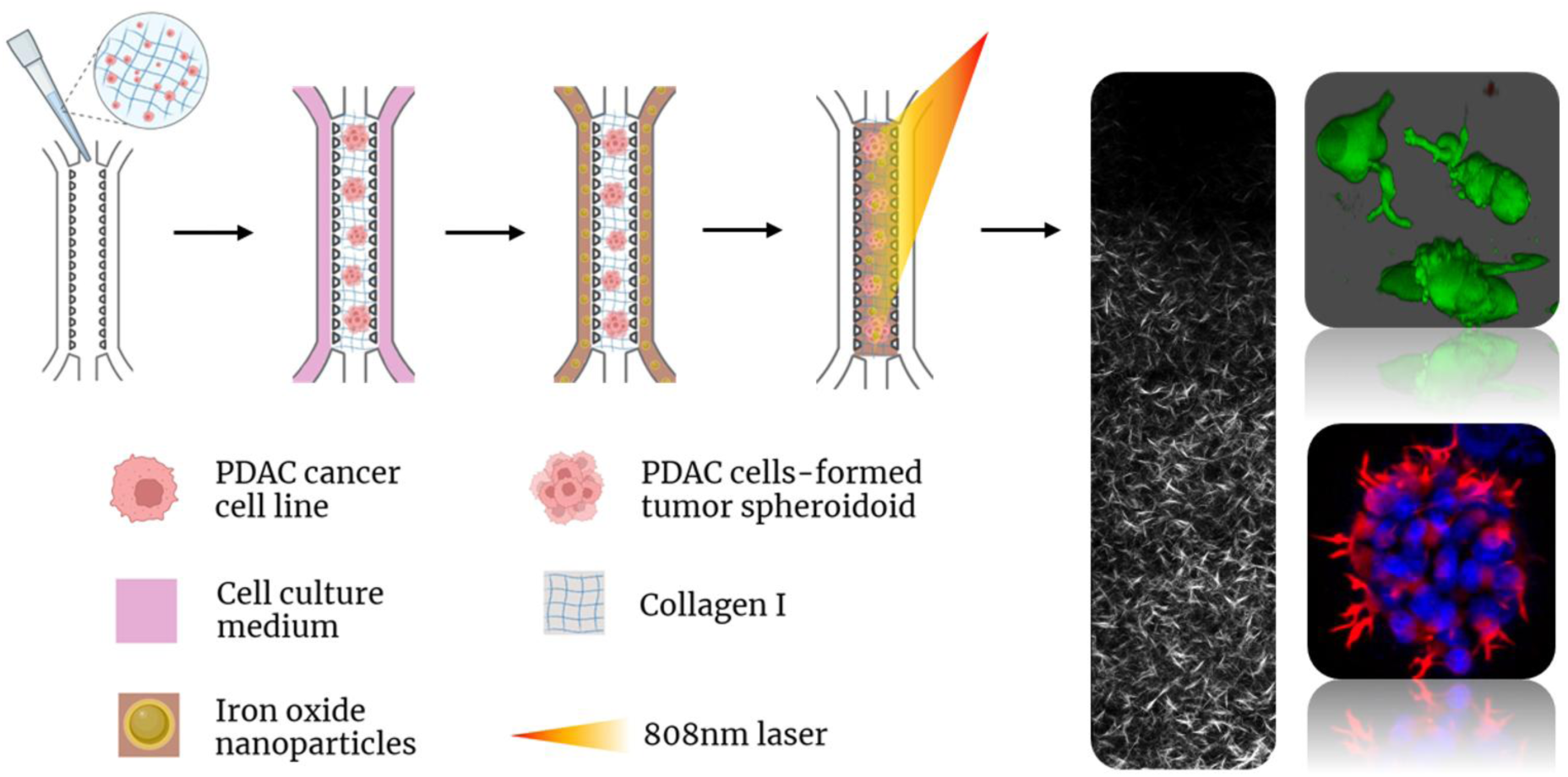
SYNOPSIS TABLE OF CONTENTS.

## INTRODUCTION

Pancreatic ductal adenocarcinoma (PDAC) is a highly aggressive tumor comprising over 90% of pancreatic cancer cases, with a 5-year survival rate of less than 8% (1,2). Its incidence and mortality rates are strongly correlated, contributing to its status as one of the deadliest cancers globally (3). The biological hallmark of PDAC is its complex microenvironment characterized by a dense tumor stroma occupying up to 90% of the tumor volume due to excessive protein secretion and deposition into the extracellular matrix (ECM). Consequently, a cascade of mechanical and biological events take place, notably hypovascularity, increased interstitial fluid pressure, hypoxia, and immunosuppression, ultimately hindering drug penetration (4–6).

Current PDAC treatments remain limited as resectable PDAC mostly rely on surgery combined with chemotherapy (7) while advanced PDAC are managed with chemotherapy only yielding limited efficacy (median overall survival: 6-12 months) (2). To face the pitfalls of current treatments, innovative approaches are currently of principal interest. An emerging tool that could address the specific features of PDAC and the limitations of its current treatments is hyperthermia. Hyperthermia involves a local temperature increase within the tumor in a range of 38-48°C with the aim of inducing a tumor-suppressing effect. Given PDAC’s significant stromal component, hyperthermia becomes yet more valuable for combination therapy development by simultaneously targeting both cancer cells and tumor stroma. Generic hyperthermia involves external irradiation modalities, such as microwaves, radiofrequency, ultrasound, global heating – all presenting challenges in localization of impact. In contrast, the introduction of nanoparticles to the field allowed for their consideration as intrabody localized heat sources that can be remotely activated to release heat locally into the surroundings. Combining hyperthermia with nanoparticles permitted precise cancer tissue targeting in both *in vitro* and *in vivo* studies with minimized side effects on healthy tissue thanks to tumor-localized impact achieved through distant stimulation (8–13). Besides, since the arrival of nanoparticles use in medicine, hyperthermia is also being regarded as an adjunctive therapy to established chemo-, immune-or radiotherapy, to increase the chemotherapeutic or ionizing radiation effects (14–18). Specifically, hyperthermia has the capacity to affect both the cancer cells and their surrounding extracellular matrix within the tumor niche giving further potential to overcome poor drug penetration limitation through synergistic effect (14–16,18). Nanoparticle-mediated magnetic hyperthermia (MHT) initially emerged as the state-of-the-art technique for achieving localized temperature elevation (8,10,12,19). However, the effectiveness and clinical relevance of MHT have been challenged by the need for high nanoparticle concentrations to induce significant heating. Plasmonic nanoparticles, such as Au/Ag nanorods, nanoshells, nanostars, were subsequently introduced for nanoparticle-mediated photothermal therapy (PTT), leveraging their efficacy as efficient light absorbents due to their localized surface plasmon resonance properties in the near-infrared (NIR) region and subsequent heat dissipation into the surrounding tissue. Remarkably, PTT demonstrates efficient performance at significantly lower nanoparticle concentrations compared to MHT (13,20,21), yet introducing another drawback, which is the limited penetration of light into deep-seated tumors. Nevertheless, PTT is now increasingly envisioned for nanoparticle-mediated anticancer thermal therapy in clinical settings. Interestingly, magnetic NPs with a magnetite core have recently been described as good competitors to the plasmonic ones for photothermal conversion due to their valence band transition between Fe^2+^ and Fe^3+^ providing absorbance in the NIR region (22,23). This, in turn, expanded the use of magnetic NPs from MHT agents to promising PTT candidates. In addition, the growing interest in magnetic NPs’ use for PTT is also driven by the innate properties of iron biocompatibility and biodegradability within tissues that leverages them over commonly used metals like gold or silver. Beyond reduced toxicity, magnetic NPs offer further possibility of employing the magnetic properties of the material, notably, as contrast agents for MRI or via magnetic guidance to the tissue of interest.

Given the recent emergence of magnetic nanoparticles for PTT as well as the yet nascent stage of PTT in clinical cancer treatment (24,25), there is now a need to fully understand the effects of these treatments, both on cancer cells and their microenvironment. To achieve this, relevant in vitro models must replicate the complexity of tumor tissues as closely as possible while maintaining precise control over experimental conditions. This is particularly important for novel cancer treatment development, as it has been demonstrated that the composition of the tumor microenvironment (TME) is a key player in the tumor response to drugs (26–29). Current strategies such as 2D models or spheroids/organoids offer fast preliminary screening but do not allow for a tunable 3D ECM, while animal models do not respond to treatment similarly to humans. In turn, tumor-on-chip models offer a promising alternative to recapitulate the 3D human TME *in vitro* while providing a fine control over the system to investigate therapeutic modes of action comprehensively (30–35). Up to date, several PDAC tumor-on-chip models have been developed (36–40), however, most of them were primarily focused on model development rather than exploring novel therapeutic strategies. While to the best of our knowledge, there have been no reports of MagNP-PTT applications in tumor-on-chip, several studies of nanoparticle-based PTT for anticancer applications have been conducted both *in vitro* and *in vivo* for some tumor types (41), including PDAC (42,43). They have demonstrated several successful results including tumor volume reduction, stimulation of antitumor immunity and effective cancer cell targeting. However, there is an evident gap for studies on nanoparticle-based PTT in tumor-on-chip platforms that we are aiming to bridge with this study.

Herein, we propose to incorporate MagNP-PTT into a microfluidic chip for investigating the impact of localized heating on the TME within a PDAC tumor-on-chip model. Specifically, this was achieved using magnetic iron oxide nanoparticles as an internal heat source within tumor-on-chip, exploiting their capacity to be stimulated with a NIR laser. The objective is to examine the response of the key components of simplified PDAC tumor-on-chip model combining pancreatic cancer cells and collagen type I-based ECM to such treatment. As primary indicators of the MagNP-PTT effect, we assessed collagen matrix denaturation and cancer spheroid cell death across a range of hyperthermia temperatures. Initially, we demonstrated a temperature-dependent sharp transition of the collagen matrix from an organized fiber mesh to the absence of defined structure around 46°C, with little-to-no detectable effect at lower temperatures. When evaluating the effect of MagNP-PTT on tumor cells, progressive cancer cell death and tumor spheroid deterioration was observed with increasing temperatures. In addition, we have identified that cancer cell viability is influenced by the duration of the rest period after MagNP-PTT treatment and before the assessment of cell survival. Extended resting periods of 24 hours have shown higher viability in samples heated at temperatures ranging from 40-44°C compared to those assessed after a 2-hour rest period. However, such an effect was not observed at higher temperatures (44-48°C). Investigating the phenotypic changes of the surviving cells we have demonstrated potential upregulation of EMT marker expression. Overall, these results establish the analysis of the MagNP-PTT –triggered impact on a tumor-on-chip and set a solid foundation for future development of the synergistic therapies of MagNP-PTT with some of the established anticancer therapies where the therapeutic window combination will play a key role.

## RESULTS

### Magnetite Nanoparticles for NIR Photothermia

Recently, iron oxide nanoparticles have emerged as efficient photothermal agents, surpassing their prior role as agents for magnetic hyperthermia. However, for effective photothermal conversion, magnetite Fe_3_O_4_ is preferred over maghemite Fe_2_O_3_. This preference arises from the mixed Fe^2+^/Fe^3+^ valence state of Fe_3_O_4_, which induces NIR optical absorbance via intervalence charge transfer (23,44,45), a feature absent in γ-Fe_2_O_3_, which only contains Fe^3+^.

It is nonetheless admitted that maintaining iron in its stochiometric +2-oxidation state in ultrasmall chemically synthesized nanoparticles (d < 10 nm) can be challenging because of the oxidation of magnetite nanoparticles to maghemite at low temperatures in water (44). Therefore, magnetic nanoparticles are generally considered to be at least partially oxidized to maghemite. Remarkably, larger biogenic magnetosomes synthesized by magnetotactic bacteria retain their magnetite composition and are often described as the only magnetic crystals allowing for pure magnetite. Magnetic nanoparticles synthesized through sol–gel microwave-assisted method was recently characterized as magnetite-composed, by Mossbauer or Fourier transform infrared spectra (46). This is exceptional for small nanoparticles with a high surface-to-volume ratio, which typically favor the formation of maghemite in aqueous media. Notably, the benzyl alcohol route not only produces highly crystalline nanoparticles but also acts as a reducing agent, converting ferric salt (iron (III) acetylacetonate) into Fe^2+^ ions. Besides, this synthesis allows for a fine control of coating agents for stabilization (47). Herein, PO-PEG-NH_2_ coating was selected (Figure 1A) to increase NPs’ stability and optimize NPs’ interactions with biological systems thanks to the hydrophilic and antifouling properties of PEG as well as positive zeta potential of the amino group (Figure S1). The synthetized nanoparticles displayed an average diameter of 9 nm as observed with TEM imaging (Figure 1B). PTT efficiency at 808 nm is attributed to the absorbance in the NIR region (Figure 1C, S2) due to the high proportion of magnetite within the as-synthesized iron oxide nanocrystals. The Mössbauer spectrum (Figure 1D) consists of asymmetrical magnetic sextet with broadened and overlapped lines. The fitting model involves considering a distribution of hyperfine fields correlated to that of isomer shift. The mean isomer values (< d> @ 0.50) which remain independent on the fitting procedure are consistent with the coexistence of magnetite and maghemite. The mean isomer shift confirms that magnetite is the dominant component, accounting for 79%. The PEG coating on the nanoparticle surface is confirmed by FTIR, showing characteristic bands at 580 cm⁻¹ for Fe-O vibrations from the iron oxide nanoparticles and at 1110 cm⁻¹ for C-O vibrations from the PEG chains (Figure 1E), indicating successful PEGylation. Consequently, they were tested for photothermal conversion in a standard tube setup (Figure 1F), using a volume of 100 µL, increasing NP concentrations (2 to 64 mM) and employing laser power densities of 1 W/cm² at 808 nm. The resultant heating (Figure 1G) positioned them as efficient NIR photothermal agents, approaching the efficacy of magnetosomes (13).

**Figure 1:**
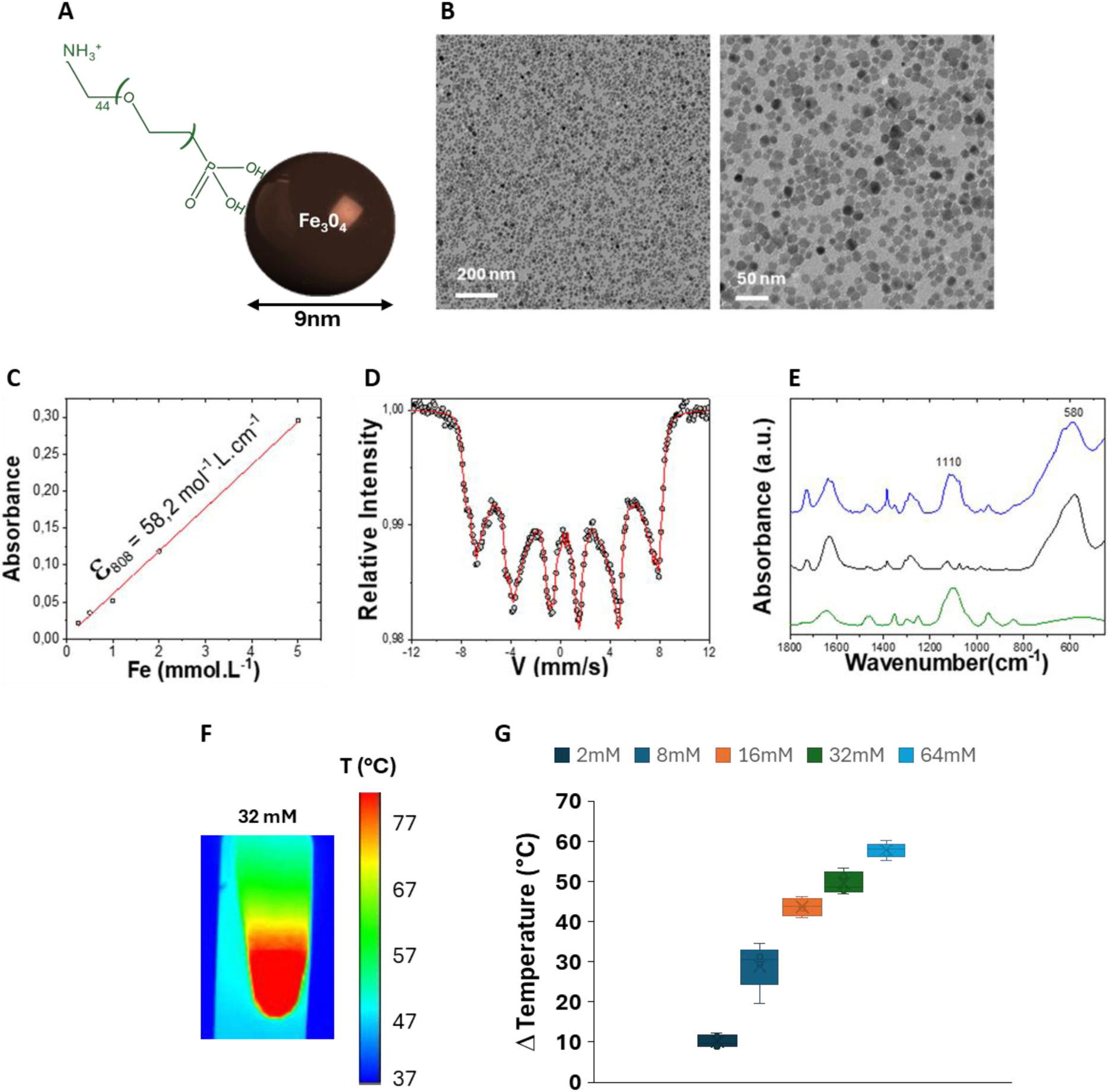
Magnetite nanoparticle physico-chemical investigation. **A**. Schematic representation of nanoparticle coating. **B.** TEM imaging of nanoparticles in suspension. **C.** 808 nm absorbance measurement as a function of iron concentration. **D.** Mössbauer spectra for the as-synthesized 9 nm NPs (black circles), with the calculated spectrum representing a mixture of 79% magnetite and 21% maghemite (red line). **E.** FTIR spectra of PO-PEG-NH_2_ molecules (green curve), bare 9 nm NP (black curve), and 9 nm NP coated with PO-PEG-NH_2_ (blue curve). **F.** Representative image of the thermal camera recording of in-tube NPs [32mM] heating with an 808nm laser at 1 W/cm^2^. **G.** Temperature profile change of in-tube NPs heating at concentrations 2-64mM and laser power 1 W/cm^2^.

### PDAC Tumor-on-Chip 3D Model

To recapitulate PDAC tumor-on-chip, a mixture of pancreatic cancer cells (PANC-1) and collagen type I (6mg/mL) was introduced into the central channel of a chip (AimBiotech^TM^, Figure 2A) enabling a capillary-based confinement of the hydrogel with cells in the central chamber (Figure 2B). This straightforward PDAC tumor-on-chip model comprising PANC-1-grown tumor spheroids in collagen type I matrix on a chip was chosen to assess the effect of MagNP-PTT treatment on the tumor microenvironment. In brief, PDAC cells were seeded in collagen I on chip and left to mature and self-organize into tumor spheroids within the matrix for a week before the MagNP-PTT exposure. After this 7-day period, cells spontaneously formed small tumor spheroids of around 100 µm in diameter (Figure 2C, D0 vs D7). The viability of these spheroids was confirmed with live/dead assay, where the green (live) signal significantly prevails over the red (dead) (Figure 2D). In addition, the morphology and volumetric distribution of spheroids within the collagen matrix was visualized via 3D confocal imaging (Figure 2E). Spheroids of different sizes were evenly distributed within the chip, with some spheroids developing protrusions (Figure 2F, Figure S4), which can be indicative of invasive cancer cell behavior. In addition, we visualized our tumor microenvironment system with PDAC spheroids uniformly surrounded by collagen fibers (Figure 2G-I). This reflects the dynamic and reciprocal interplay between the tumor spheroids and their microenvironment, which is also highly characteristic of the *in vivo* tumor niche (29,48).

**Figure 2:**
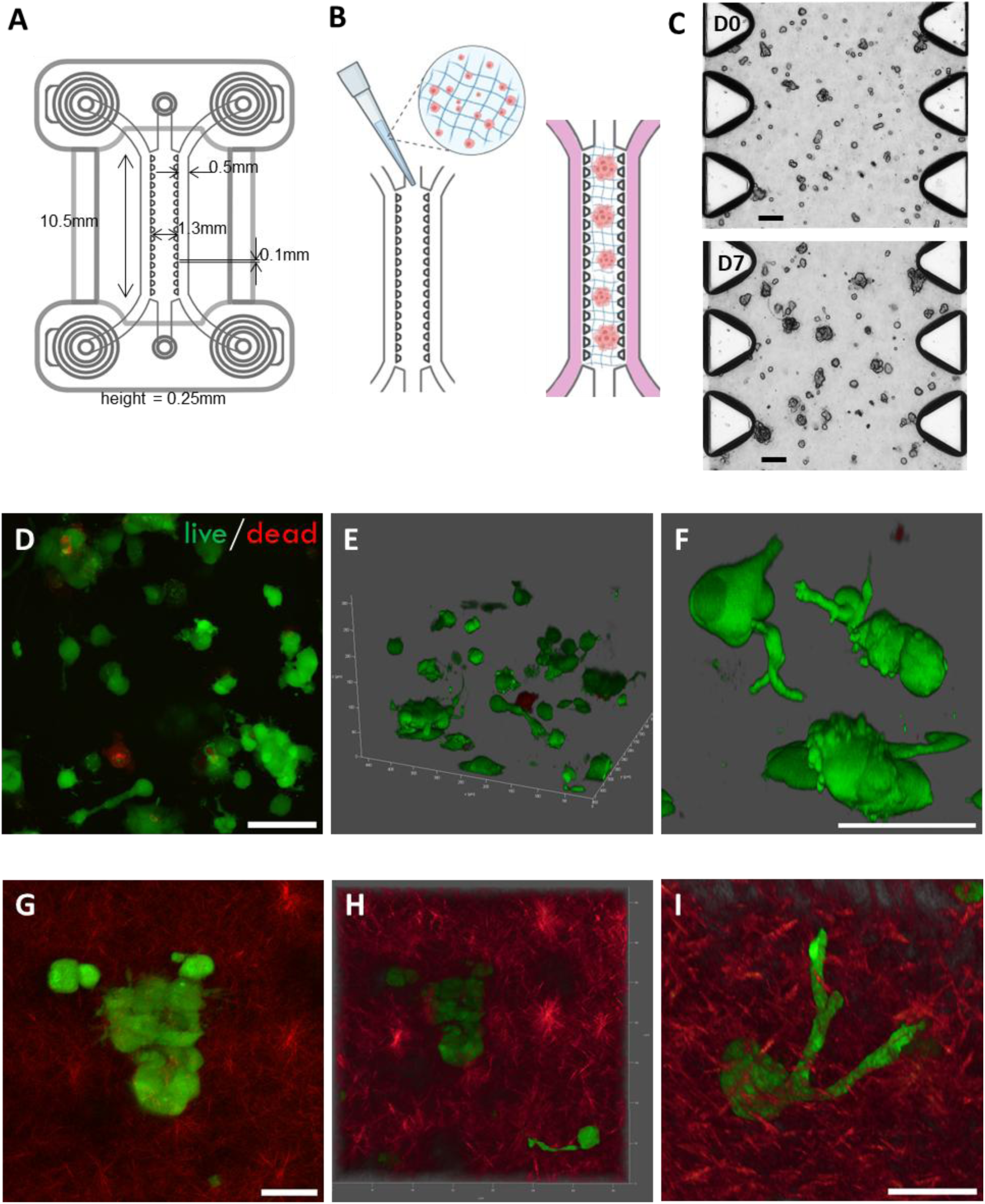
Development of the 3D PDAC tumor-on-chip model. **A**. Schematic representation of microfluidic chip idenTx3, AimBiotech^TM^. **B.** Schematic representation of cancer cell seeding in collagen on chip with subsequent spheroid formation. **C.** Bright field microscopy images of PDAC tumor spheroid development from single cells (D0) vs after 7 days of maturation in collagen on the chip (D7). Scale bar = 100µm. **D-F.** Confocal microscopy images of Live/Dead assay (dead = red; live = green) with (D) z-projection of the 3D stack, at day 8 in-chip, (E-F) 3D confocal stack reconstructions. Scale bar = 100µm. **G-I.** Second harmonic generation (SHG) microscopy image of a tumor spheroid (green) with the surrounding collagen matrix (red) z-projection of the 3D stack (G), reconstructed 3D images (H-I). Scale bar = 50µm.

### Administration of Magnetite Nanoparticles in Tumor-on-Chip

Using the PDAC tumor-on-chip model, where cancer cells have self-organized within the matrix over one week and display reflections of a tumor-like organization, the next step involved the injection of NIR-heat-generating magnetite nanoparticles into the chip. The principle of introducing NPs into the chip (Figure 3A) involves their passive diffusion over a 24-hour period from the lateral channels, each being injected with a given NPs concentration, into the PDAC tumor microenvironment modelled in the central chamber of the chip. Figure 3B displays intact tumor spheroids in collagen on chip after 24h incubation with NPs at [Fe]=32mM, while Figure 3C illustrates the viability of these spheroids through live/dead confocal imaging 3D reconstruction, confirming the absence of adverse cytotoxic effects induced solely by the NPs. Transmission electron microscopy (TEM) was further used to localize the NPs at the nanoscale within the chip. NPs were consistently found adsorbed on the collagen fibers (Figure 3D). They were also detected both at the outer region of cells, near the plasma membrane within the ECM, as well as being endocytosed by the cells (Figure 3E). Subsequently, many NPs were identified inside endosomes within the cancer cell cytoplasm (Figure 3F). Overall, this imaging demonstrates excellent penetrability of NPs within the collagen-based matrix, successful NP endosomal internalization by tumor spheroid cells and lysosomal storage confirming their ubiquitous presence in our system. Additionally, Perls Prussian Blue staining performed *in situ* within the chips detected the presence of iron in tumor-on-chip injected with NPs as compared to control, with blue coloration observed inside and around the spheroids as well as diffused within the matrix (Figure 3G). The intensity of the blue color increased with NP concentration, 8 vs 32 mM, and blue extracellular matrix coloration surrounding the spheroids was most pronounced at 32 mM.

**Figure 3:**
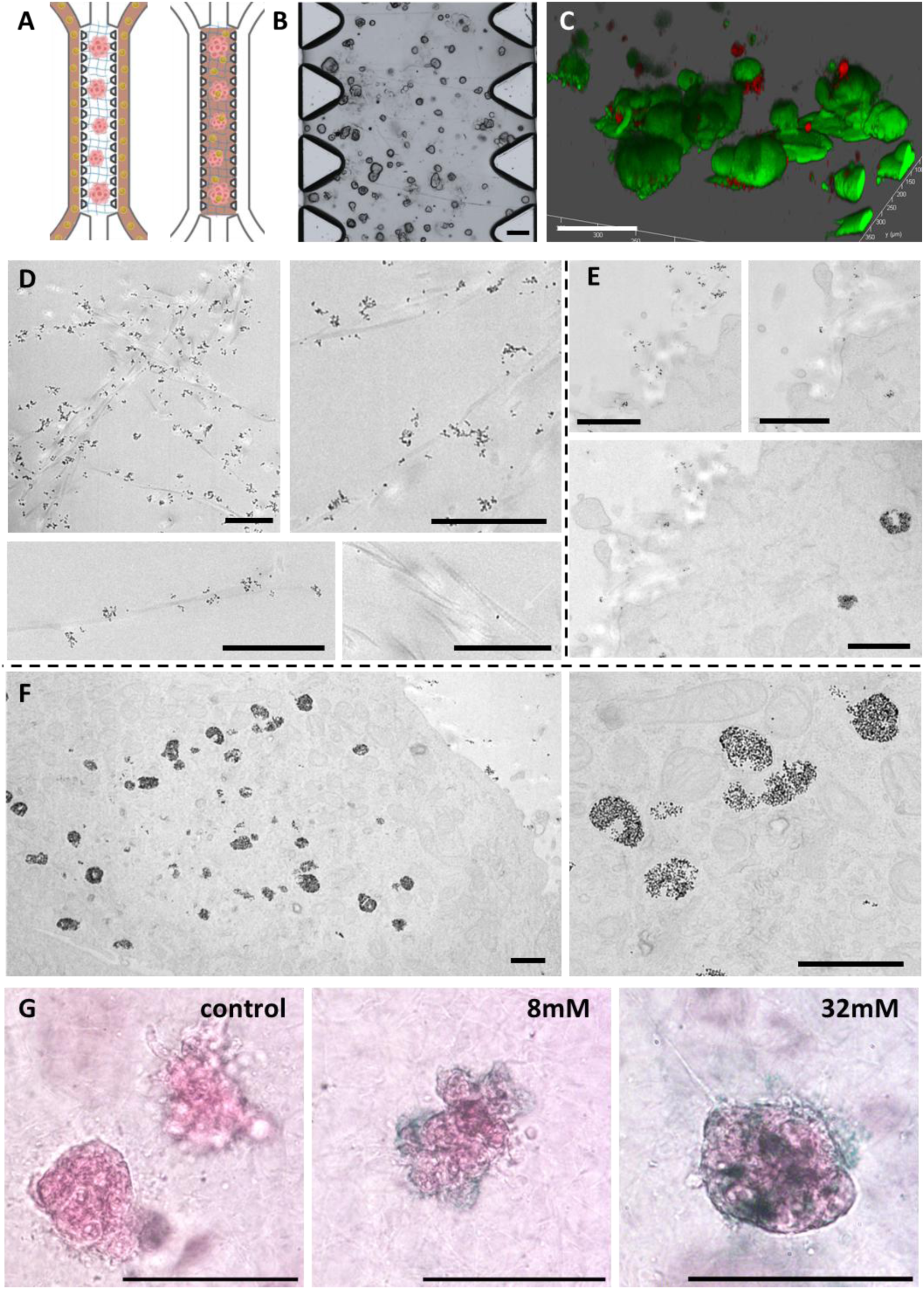
Administration of the magnetite nanoparticles and imaging within tumor-on-chip. **A.** Schematic representation of on-chip NP administration and diffusion. **B.** Bright field microscopy image of tumor spheroids on chip on D8, 24h after NP incubation. **C.** Confocal microscopy images of Live/Dead assay (dead = red; live = green), 3D confocal stack reconstruction of tumor spheroids on chip on D8, 24h after NP incubation. Scale bar = 100µm. **D-F.** TEM imaging of NPs adsorbed on collagen fibers (D), NPs endocytosis in cells (E), NPs internalized on cell endosomes (F). NPs appear as dark contrast. Scale bar = 1µm. **G.** Perls Prussian Blue staining of iron deposits (in blue) in PDAC spheroids in collagen on chip. Scale bar = 100µm.

### Implementation of MagNP-PTT in PDAC tumor-on-chip and its Impact on Collagen Matrix

The next step was to irradiate the NP-containing chips with an 808 nm NIR laser. A dedicated setup was developed to enable irradiation of microfluidic chips with a laser and measure the induced temperature elevation (Figure 4A). The setup consists of a water bath-supplied holder for the chip, which maintains microenvironment within the chip at 37°C, an 808nm laser unit, and an infrared thermal camera to visualize and record temperature change. The optimal distance between the laser and the chip was identified to be 6.1cm (Figure S3) to maximize the homogeneity of the surface laser power distribution (i.e., flatter laser power Gaussian distribution) while maintaining high attainable laser power range. This configuration helped reduce, though not completely eliminate, temperature variation along the length of the chip during laser exposure.

**Figure 4:**
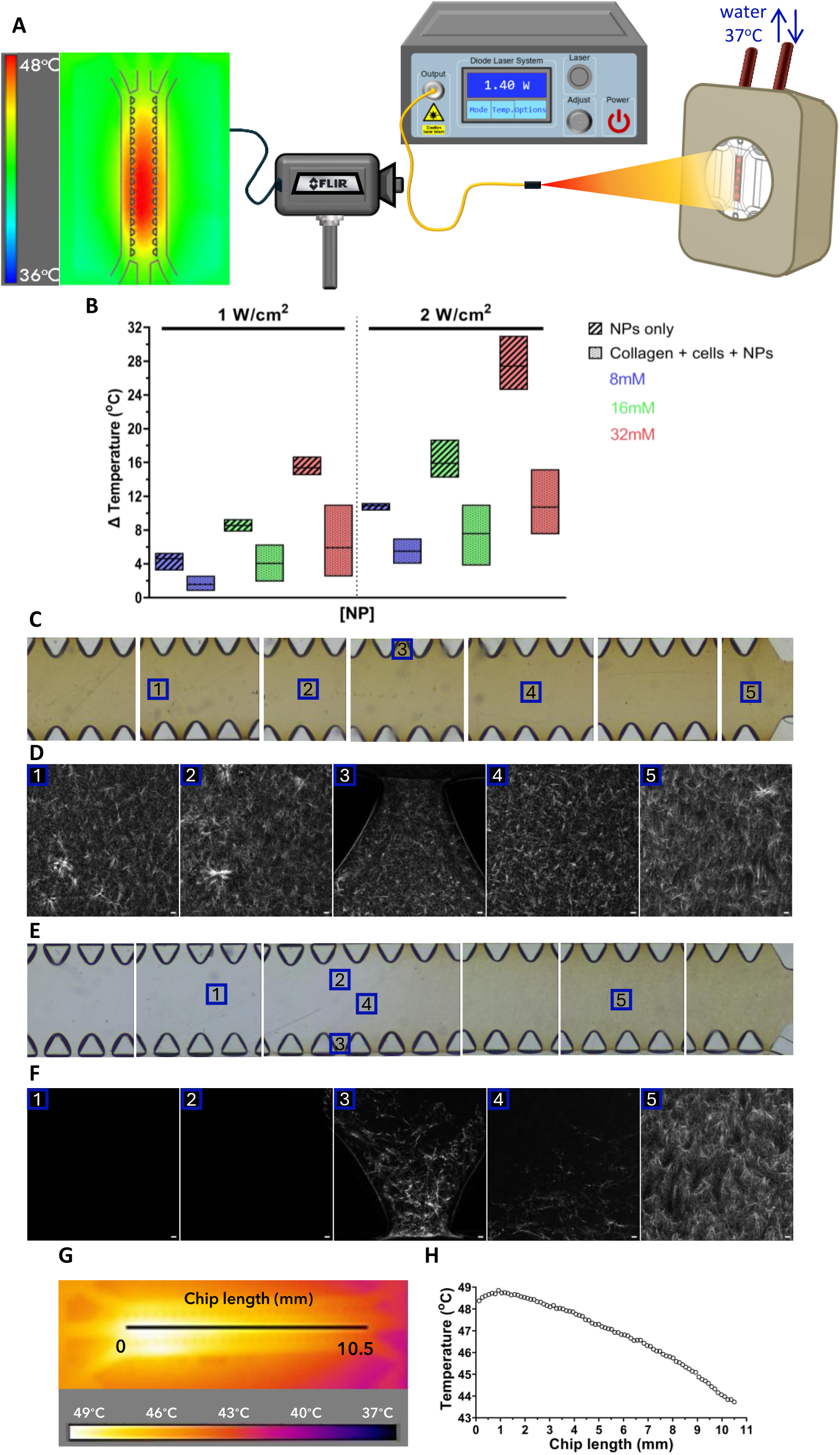
Nanoparticle-mediated photothermia heating setup and matrix impact characterization. **A.** Scheme of MagNP-PTT setup. **B.** Temperature increase (ΔT) as a function of NP concentration and surface laser power for NPs heating in chip VS NPs diffused in collagen + cells on chip. Data are represented as a boxplot with line at mean. **C.** Bright field images of the control chip with NPs at 32mM diffused in collagen. **D.** Collagen hybridizing peptide staining imaging of different areas in the control chip with 32mM NPs with confocal microscopy. **E.** Bright field images of the chip (16mM NPs) with partially denaturated collagen. **F.** Collagen hybridizing peptide staining imaging of variously denaturated areas in the exposed chip with 16mM NPs, confocal microscopy. **G.** Corresponding thermal camera heating recording within the chip with representative temperature gradient along the length of the chip. **H.** Temperature variation as a function of the chip length.

We next characterized our MagNP-PTT setup by investigating both the temperature increase in the presence of NPs as well as its distribution within the system. The initial step was to assess the efficiency of light-to-heat conversion in the system design. NPs were introduced into the central channel of the chip either via direct injection of the nanoparticle solution (in water) or through their diffusion over 24-hour period from the lateral channels into the collagen type I (6 mg/mL) matrix containing pancreatic cancer spheroids. Temperature elevation was measured for increasing iron concentrations of NPs (8, 16, and 32 mM) and laser power densities (1 and 2 W/cm^2^). For both conditions (NPs in water and diffused in the collagen with embedded cells), a similar pattern of temperature increase was observed with increasing NP concentration and laser power (Figure 4B). However, the overall temperature increase was higher for NPs heating in water compared to NPs within collagen with cells. This suggests reduced NP diffusion within collagen matrix, potentially due to the interactions of NPs with collagen fibers, as observed by transmission electron microscopy imaging (Figure 3D). As a result, with the given NP concentration and laser power ranges we can elevate the temperature within the tumor-on-chip by ΔT=1-12°C (starting from 37°C) and thus cover the full range hyperthermia temperatures (38-48°C) in the tumor-on-chip.

To further evaluate the effect of MagNP-PTT on tumor-on-chip, the next step involved investigating how MagNP-PTT-induced heating affects the collagen type I matrix. For this, collagen matrix at 6mg/mL on chip was subjected to NP-mediated laser heating and subsequent collagen fiber imaging to examine structural impact. While second harmonic generation (SHG) microscopy is a state-of-the-art method for collagen fibers visualization, collagen hybridizing peptide (CHP) staining is emerging as well due to its ability to exclusively bind to collagen fibers through triple helix hybridization (49). To explore the comparability of the two techniques, we first imaged untreated collagen matrix on chip (control) with two-photon microscopy to simultaneously observe the SHG and CHP signals (Figure S5). As a result, both images revealed an isotropic mesh of fibrils of around 1 µm in diameter. Remarkably, the SHG signal was fully overlapped by the CHP staining signal, demonstrating excellent comparability on methods’ output. However, the CHP fluorescent image showed more signal present than the SHG image, suggesting that CHP staining provides with a higher degree of sensitivity, allowing to visualize collagen degradation at microstructural level (49–51).

MagNP-PTT treatment was then applied to assess its impact on collagen matrix structure. While several studies have investigated collagen I thermal denaturation under various temperature modulation approaches as well as analytical methods (52–54), results show variability in collagen denaturation points and responses to increased temperature. Factors influencing this variability include collagen source, extraction method, concentration, solvent, degree and/or method of crosslinking, among the others (55). For type I collagen, the range of temperatures leading to irreversible thermal changes have been reported to be between 37°C and 55°C, where the loss of fibrillar structure was observed (56).

In this study, to assess the impact of NP-mediated heating within our collagen matrix on chip, NPs were let to diffuse from the lateral channels of the chip into the collagen matrix for 24 hours before heating, followed by the CHP staining imaging to study collagen structure changes after laser exposure. The control chip (Figure 4C) with 32mM NPs diffused in a chip demonstrated no significant matrix structure changes across the chip (Figure 4D) Chips, containing NPs at 16 mM (Figure 4E) and 32 mM (Figure S6) concentrations, were exposed to the laser powers at 2.5 W/cm^2^ and 1 W/cm^2^ respectively for 20 min, heating the matrix within the range of temperatures known to cause collagen type I degradation. The resulting temperature gradient along the chip ranged from 44°C to 49°C for 16mM (Figure 4 G-H) and from 42°C to 48.5°C for NP at 32 mM (Figure S6 E-F). These gradients induced varying responses within the collagen matrix, allowing not only to explore the impact of a wider range of hyperthermia temperatures within a unique matrix but also represents a better model of temperature distribution of the *in vivo* setting. We observed a rather dichotomous effect, where no significant matrix alterations were detected at temperatures below 46°C with a drastic disappearance of detectable collagen fibers around 46°C, consistent with previous reports for similar collagen source (56). Bright field images of the chip revealed transparent zones where temperature increased above 46°C, while areas at lower temperatures remained light brown due to the presence of NPs in the collagen (Figure 4E). CHP staining confocal images corroborated these observations, showing a transition from intact collagen matrix with a dense network fibril network (Figure 4F [5]), similar to the control (Figure 4D), to nearly fully denatured fibrils that were no longer detectable (Figure 4F [1-4]). Depending on the temperature gradient across the sample centered around the collagen denaturation point, an aperture within the collagen matrix could form (Figure S6 C) with different fibril organizations around it. Two outcomes upon partial collagen matrix denaturation were observed: i) a gradual matrix transition from a well-defined fibril mesh to the absence of visible fibrils (Figure S6 D [2-4]) and ii) accumulation of detected collagen damage at the aperture border (increased CHP fluorescence, Figure S6 D [1]), likely, due to collagen fiber accumulation from matrix folding caused by contact point loss after denaturation. Therefore, although there is a clear transition from an intact to an undetectable matrix state due to fibrillar denaturation, a more complex matrix restructuring occurs at a finer level around the denaturation point.

### PTT-Induced Cell Death in PDAC tumor-on-chip: Effects of Rest Time and EMT Impact

Following the MagNP-PTT set up for tumor-on-chip applications and PDAC tumor-on-chip model establishment, we next studied the effect of local heating of on-chip PTT on PDAC tumor spheroid survival. The combination of different NP concentrations (8, 16 & 32 mM) and surface laser powers (1-2 W/cm^2^), as illustrated in Figure 5A, allowed to heat the samples in the overall range of 38 to 48°C. Figure 5A also showed that once the laser is turned on, the temperature in the chip rises during the first few minutes followed by a plateau that remains until the laser is off. The cell death analysis was then divided into 5 temperature study groups stemming from the final temperature reached within the sample, grouped into five 2°C-range bins starting from 38 to 48°C. Figure 5A evidenced that all the intra-group temperature variations in time are well confined by the predefined 2°C temperature window for all the groups. Besides, temperature profiles were extracted as a function of position in the width of the chip (Figure 5B) across all the samples used in this study. Temperature distribution in the width of the chip, more specifically the central chamber enclosed by the pillars, is homogeneous within each group and consistent between the groups. A respective control with PTT exposure at 2 W/cm^2^ without NPs was performed to confirm the absence of heating and negative effect on the tumor spheroids in the absence of NPs in the system (Figure S8).

**Figure 5:**
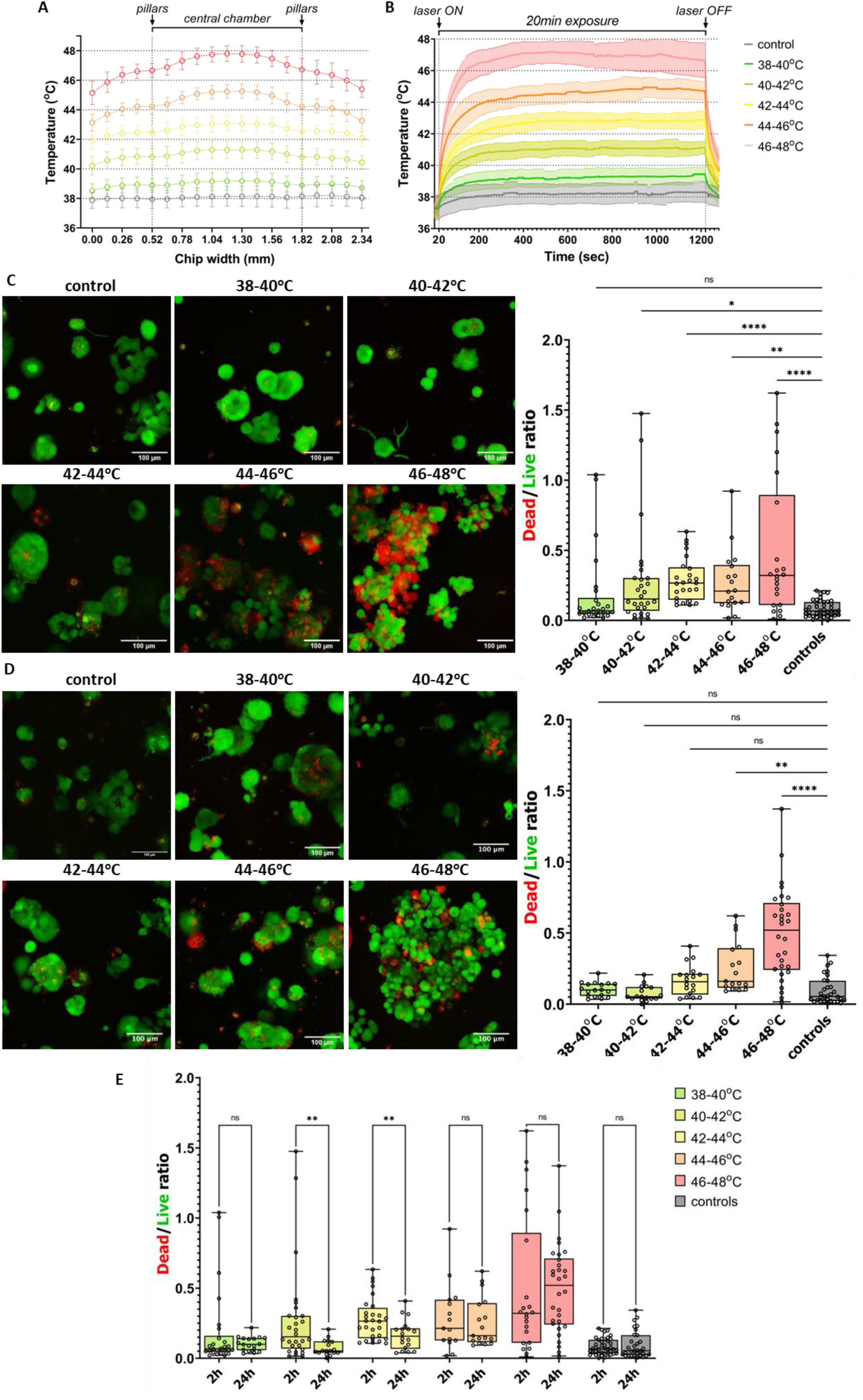
Temperature-dependent cells death evaluation after MagNP-PTT. **A**. Temporal temperature profile within the chip over 20 min of laser exposure for the 5 studied temperature groups. Data are represented as mean ± SD of all the experimental data (n>3). **B**. Spatial temperature distribution profile along the width of the chip, including central channel with two side channels, for the 5 temperature groups studied. Data are represented as mean ± SD of all the experimental data (n>3). **C.** PDAC cell viability in collagen on chip 2h after MagNP-PTT exposure. Representative confocal microscopy images (Z projection, SD type) for each of the studied temperature groups, live/dead (Calcein Green/Texas Red) viability assay. Scale bar = 100µm. Live/dead assay quantification showing cell death estimate as a function of the temperature group, 2h rest time after MagNP-PTT exposure. **D.** PDAC cell viability in collagen on chip 24h after MagNP-PTT exposure. Representative confocal microscopy images (Z projection, SD type) for each of the studied temperature groups, live/dead (Calcein Green/Texas Red) viability assay. Scale bar = 100µm. Live/dead assay quantification showing cell death estimate as a function of the temperature group, 24h rest time after MagNP-PTT exposure. **E**. PDAC cell recovery evaluation after MagNP-PTT exposure. Comparison of dead-to-live signal ratio for 2h vs 24h after laser exposure for all the studied temperature groups as an estimate for cell recovery after temperature increase. Data are presented as box & whiskers plot (min to max) including all experimental data points (>3 independent experiments for each condition, ≥2 chips per experiment, ≥3 independent ROIs analyzed per chip). *p< 0,1; **p< 0,01; ***p< 0,001; ****p< 0,0001.

Laser exposure was performed for 20 min, and heating effects on cancer cell death were first evaluated 2 hours post-treatment (Figure 5C). For temperatures up to 40°C, the morphology and dead/live signal proportion in images was similar to the control conditions, with PANC-1 cells maintaining as compact spheroids, indicating minimal adverse effects. In contrast, tumor-on-chip systems heated above 42°C showed a progressive impact on cell viability and spheroid integrity (Figure 5C). The dead cell (red) signal gradually increased from the 42-44°C group to the two highest temperature groups (44-46°C and 46-48°C), where spheroid integrity was strongly impaired, with single cancer cells becoming more distinct (Figure 5C). To quantitatively analyze cell viability, an algorithm was developed, where the live and dead signals were segmented independently using machine learning for each condition and sample. These two signals were then quantified by computing their representative areas in 3D to determine their total presence within the sample and calculate their ratio as an indicator of cell viability. This quantitative analysis confirmed the overall trend of increasing cell death with increasing temperature (Figure 5C, right panel). Little-to-no detrimental effect was observed after heating 1-3°C above 37°C with the dead-to-live ratio median at 0.15 and the interquartile range spanning from 0.05 to 0.3. In contrast, exposure to temperatures above this range significantly impacted cell viability compared to controls. However, even in the highest temperature group, the median dead/live ratio reached was around 0.3 with the interquartile range spanning up to 0.9, indicating equal levels of dead and live cells. Thus, while MagNP-PTT caused substantial cell death due to local heating, a significant proportion of pancreatic cancer cells remained alive even after heating to 48°C for 20 min. Next step consisted in assessing the effect of rest time on cell survival after MagNP-PTT treatment. Cells were thus allowed to rest for 24 hours before evaluating their survival across the same temperature range. Confocal imaging revealed that tumor-on-chip heated at the lowest temperatures (38-44°C) exhibited high cell viability and spheroid integrity similar to controls (Figures 5D, top row). In contrast, both the second highest (44-46°C) and the highest (46-48°C) temperature groups showed significant cell death showed marked elevation of the amount of dead signal, with a noticeable increase in spheroid fragmentation into single cells. The quantitative analysis of cell death confirmed these observations, showing that tumor spheroids heated on-chip up to 44°C displayed no adverse viability effects compared to controls, while significant dead signal was observed in chips heated at 44-46°C, with even greater levels in the 46-48°C range (Figure 5D, right panel). Such study of two post-treatment times allowed to explore whether MagNP-PTT heating could induce the stratification of heated cells into three groups: dead cells from temperature increase, unaffected live cells, and damaged cells not yet irreversibly changed. This implies that over longer periods, the dead-to-live ratio may be influenced by proliferating live cells or recovering damaged cells. To assess such potential response stratification, we proceeded with the group-by-group comparison temperature range effect for 2h vs 24h post-treatment rest time (figure 5E). While no significant differences were observed for the lowest range (38-40°C) due to low cytotoxic impact of PTT, for the 40-42°C and 42-44°C groups, a marked decrease in the dead-to-live ratio from 2 to 24 hours suggests that cell proliferation and recovery were more prominent than further cell death. In contrast, for the 44-46°C and 46-48°C ranges, dead/live ratios did not differ significantly between 2 and 24 hours, remaining more elevated as compared to controls for both time points (Figure 5E). Thus, while 24 hours post-treatment may promote live cell proliferation and recovery in the 40-44°C range, these effects do not offset irreversible damage at higher temperatures (44-48°C).

Finally, a more in-depth study on cellular effects was conducted by retrieving the cells after 24 hours post-treatment with MagNP-PTT (40-44°C – temperature, at which significant recovery was observed 24h after exposure) and analyzing their gene expression. Genes involved in apoptosis inhibition (MCL1, Myeloid Celleukemia 1 and BIRC5, baculoviral IAP repeat containing) along with mesenchymal (*vimentin*, N-cadherin (*cdh2)*, focal adhesion kinase (*fak)*, fibrillin (*fbn1*), zinc finger transcription factor (*zeb*)) and epithelial markers (E-cadherin (*cdh1*), *a* –catenin (*a-cat*)) were studied. It is worth noting that detecting effects on cell death under these conditions is challenging, as the RNA being analyzed corresponds to living cells or those in recovery. As a result, apoptosis-related genes birc5 and mcl1 showed no significant changes due to the treatment suggesting that MagNP-PTT does trigger apoptosis resistance under our exposure conditions. In contrast, a moderate increase in *vimentin* and *fak* genes was observed, along with non-significant increases in *zeb* and *fbn1*. Notably, the cadherin markers (E and N), *cdh1* and *cdh2*, as well as α-catenin, remained unchanged. Thus, the mesenchymal markers (*vimentin, fak, zeb, fbn1*) increased post-treatment, while the epithelial markers (*cdh1, a-cat*) remained unchanged, indicating a slight potential for epithelial-to-mesenchymal transition triggered by the PTT treatment. In addition, this mild effect was confirmed by immunofluorescence imaging of E cadherin and vimentin on-chip, where PTT slightly increased vimentin expression in the sample post MagNP-PTT exposure (Figure 6D) as compared to controls without (Figure 6B) and with NPs (Figure 6C) that were not exposed to the laser. On the other hand, E-cadherin expression remains unchanged across samples. This may be stemming from the fact that PANC-1 are characterized to belong to a quasi-mesenchymal PDAC cell line subtype, expressing lower levels of epithelial-associated genes (57,58). All in all, these results indicate that while the surviving cells post-MagNP-PTT exposure may undergo potential EMT profile changes as evidenced by modest changes in some gene expressions, they also open a promising avenue for exploring the intricate interplay between PTT anticancer therapies and cellular plasticity. In particular, understanding these mechanisms could not only inform the optimization of thermal therapies (e.g., duration, dosage, therapeutic window etc.) but also shed light on potential synergistic strategies development with other treatments (e.g., chemotherapy, immunotherapy etc.) to harness the efficacy of cumulative therapeutic effect.

**Figure 6:**
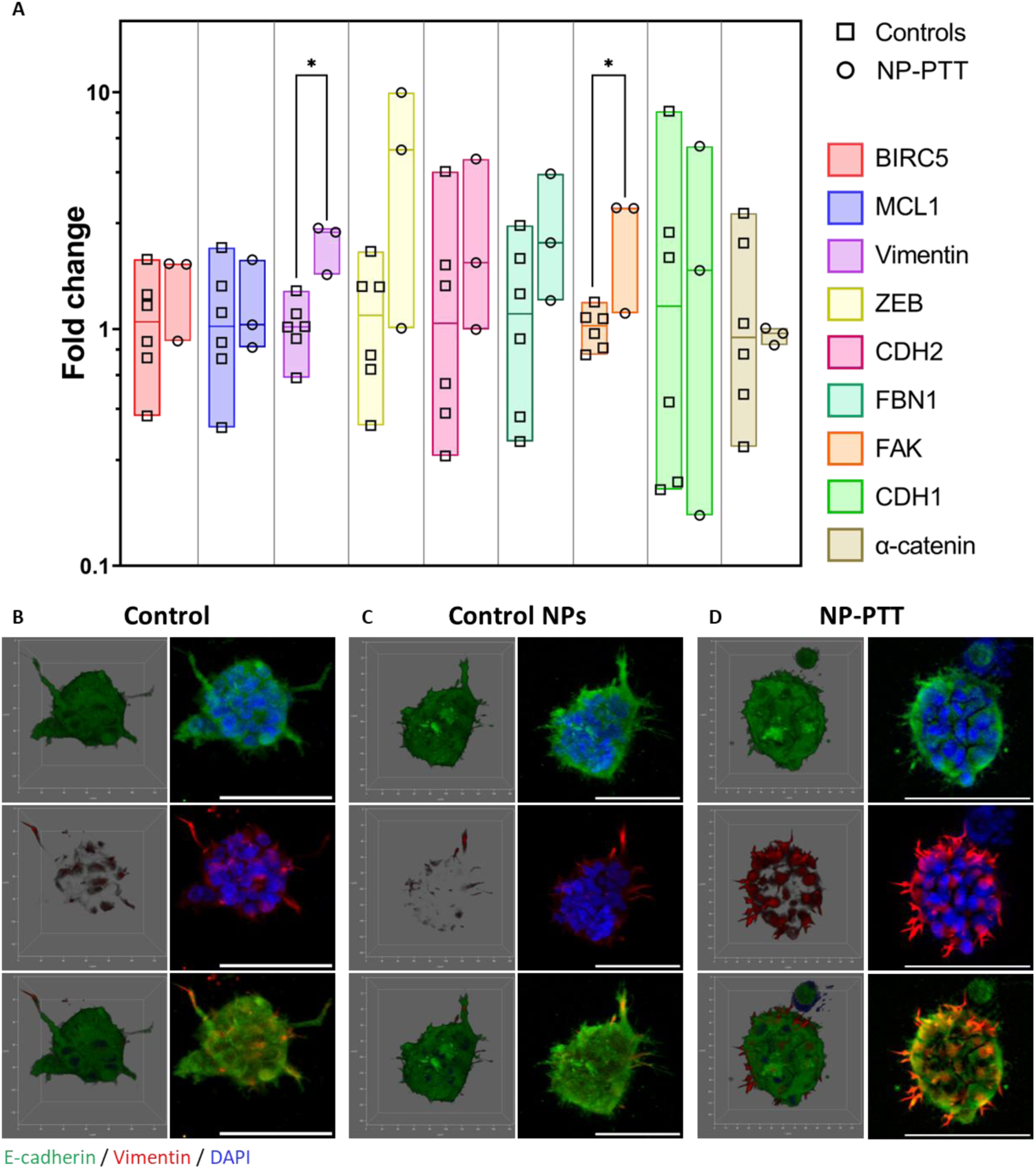
Characterization of MagNP-PTT effects of tumor spheroids gene expression. **A.** Expression of a panel of mesenchymal, epithelial and apoptotic genes measured by RT-qPCR for control samples cultured in collagen on chip and after MagNP-PTT exposure at 40-44°C, 24h rest time. **B**. 3D reconstructed confocal images (left panel) and Z-projections of 3D stacks (STD method) (right panel) of single spheroids for control, control with NPs and post-MagNP-PTT samples immunostained for E-cadherin (green), Vimentin (red) and DAPI (blue).

## DISCUSSION

With the emergence of nanotechnology for novel therapy development, the possibilities for innovative PDAC treatments expanded a lot. However, most of those nanomaterial-based strategies are mainly applied for anticancer agents’ delivery (59). In contrast, treatments with nanoparticles being the active players are still viewed under the paradigm of non-traditional, pilot approaches. Among them, magnetic hyperthermia and in particular nanoparticle-mediated photothermia have been moderately explored in the scope of anticancer PDAC treatment (42,60–64). With the recent development of magnetic nanoparticles for PTT applications, their integration into clinical cancer treatment remains at a very early stage. It is thus essential to gain a comprehensive understanding of the impacts of these therapies on both cancer cells and their surrounding microenvironment.

Our study illustrates the effectiveness of MagNP-PTT in inducing cell death within a PDAC tumor-on-chip model. The magnetite nanoparticles used, initially reported as magnetic hyperthermia agents, demonstrated efficient near-infrared light-to-heat conversion efficiency compatible with the chip design. Their ability to diffuse through the collagen matrix and penetrate cancer spheroids allowed for precise temperature control. By optimizing nanoparticle concentration and laser dose, temperature elevations could be finely adjusted, spanning from 38 to 48°C, encompassing the range of hyperthermal treatments. Thanks to our unique NP-PPT set up compatible with tumor-on-chip, we explored the impact of magnetite NP-mediated PTT both on pancreatic cancer cells and collagen type I-based ECM through heating-induced cell death as well as matrix denaturation respectively. We aimed to provide here the first example of such therapy efficiency with tumor-on-chip models with a promising outlook for *in vivo* applications.

Significant impact on tumor cell viability can be observed shortly after the treatment when heating at as low as 40-42°C and progressing to up to 48°C. This decreased cell survival is associated with spheroid disintegration and appearance of single cell aggregates suggesting the presence of adverse, potentially irreversible events. On the other hand, the amount of time cancer cells were left to rest post-treatment before evaluating their viability considerably affects their survival. Following 24h rest period, the viability of cells heated at 40-44°C was markedly increased as compared to 2h rest, suggesting the events of either residual live cells’ proliferation and/or reversibility of cell damage induced over the 24h period. In contrast, exposure to higher temperatures resulted in irreversible cellular damage certainly due to stronger, thus, more persistent adverse effects on cancer cells. It is worth noting that, although MagNP-PTT has shown a positive effect on pancreatic cancer cell death, partial cell survival persists after treatment even at high temperature. This, in turn, links to the novel paradigm of hyperthermia applications, where they are mostly considered as promising candidates for synergistic anticancer treatment strategies (65,66).

Several studies of hyperthermia combined with conventional chemotherapy have been proven successful both *in vitro* and *in vivo*, mainly through improved drug delivery (67,68). Hence, an efficient use of hyperthermia-based combination therapies may positively affect the therapeutic window of established drugs and allow to increase the efficiency of existing anticancer approaches for PDAC. Reported improvement in chemotherapy delivery is, in part, considered to occur thanks to partial ECM denaturation. These include the ECM stiffness, density, interstitial pressure among the others that are initially increased through PDAC desmoplasia-triggered chain of events – all in all leading to decreased drug penetration. Our study highlights the potential of MagNP-PTT not only for inducing targeted cell death but also for addressing the critical challenge of drug delivery in PDAC. On-chip PTT at a temperature above 46°C was found to disrupt the ECM with no detectable fibril structure. This disruption may potentially enhance drug delivery by modifying the tumor microenvironment as well as increase vascular permeability and reduce interstitial fluid pressure, overcoming the mechanical barriers posed by the stroma. Consequently, PTT not only directly targets cancer cells but might also foster a more favorable environment for therapeutic agents to penetrate effectively, reinforcing its role as a synergistic strategy.

Finally, the temperature– and time-dependent effects on cell viability and morphology underscore the importance of optimizing treatment parameters, which could improve outcomes in thermal therapies. While MagNP-PTT shows promise, translating these findings to *in vivo* models and clinical settings remains a challenge, necessitating further exploration of heat generation in a real tumor, and the long-term effects and safety of such treatments.

Overall, this work positions MagNP-PTT as a compelling strategy for treating PDAC, paving the way for more effective therapeutic interventions. The precision of NP-mediated PTT offers solutions for targeted therapy, with the capability to selectively induce cell death in tumor spheroids while minimizing damage to surrounding healthy tissue. Furthermore, the tumor-on-chip model used in this study provides valuable insights into the complex interactions between nanoparticles, the ECM, and cancer cells, enhancing our understanding of tumor dynamics and guiding the development of more effective therapeutic strategies. Future studies could focus on the intricate interactions within the tumor microenvironment by developing advanced PDAC tumor-on-chip model to include more of the crucial players in PDAC establishment and progression, such as pancreatic stellate cells and other ECM proteins, and the potential integration of MagNP-PTT with current chemotherapeutic treatments to enhance patient outcomes.

## CONCLUSION

In conclusion, this study presents the implementation of NP-mediated photothermal therapy for tumor-on-chip applications using a simple PDAC tumor-on-chip model. Evaluation of MagNP-PTT effect on the PDAC tumor microenvironment showed promising results in inducing targeted, temperature-dependent cancer cell death and collagen extracellular matrix degradation. Furthermore, gene expression analysis revealed a modest increase in mesenchymal markers following PTT, while epithelial markers remained unchanged, indicating a mild epithelial-mesenchymal transition. These findings underscore the potential of MagNP-PTT therapy while setting the foundation and underlining the importance of further investigation and optimization of MagNP-PTT treatments for potential *in vivo* applications. Moving forward, further studies into the responses of cancer cells and stromal components to MagNP-PTT in a more complex PDAC tumor-on-chip model, integrating pancreatic stellate cells and ECM proteins, such hyaluronic acid, will be essential for understanding the broader impacts of MagNP-PTT on pancreatic cancer.

## METHODS

### Cell line and culture

In this work, human pancreatic cancer cell lines PANC-1 were used, purchased from the American Type Culture Collection (ATCC, CRL-1469TM). Cells were continuously cultured in 2D in T75 flasks in a humidified incubator at 37°C with 5% CO_2_ in high glucose DMEM medium (Gibco 61965-026, GlutaMAX^TM^) supplemented with 1% of penicillin-streptomycin (P/S) (Gibco 15140-122), and 10% fetal bovine serum (FBS) (Gibco 10270-106), with each passage taking place at 80-90% confluence level and proceeding from passage 3 to 20.

### 3D culture on chip

Commercially available microfluidic devices (DAX-1, idenTx 3, AIM Biotech, Singapore) suited for 3D cell culture were used in this project (gas-permeable cyclic olefin copolymer (COC)-based material with central chamber width 1.3mm, side chamber width 0.5mm, height 0.25mm, pillar distance 0.1mm). They were cut to obtain single chips from the initial plate of 3 and sterilized with UV before usage.

To model the PDAC microenvironment on chip, PANC-1 single cells were seeded in collagen type I in the central chamber of the chip. For collagen mixture, collagen type I HC, rat tail (Corning, USA, ref. 354249) was mixed with PBS 10X, NaOH 1M, MiliQ H_2_O according to manufacturer’s instructions to yield 6mg/mL mixture of physiological osmolarity and pH. The cell seeding suspension was prepared by detaching cells from the flask at 80-90% confluence and resuspending in the collagen mixture. The cell-collagen suspension was then introduced in the central chamber of the chip at concentration of 500 000 cells/mL (∼5000 cells/chip). All the manipulations with collagen were performed at 4°C to avoid premature collagen polymerization. After seeding, collagen with cells in the chip is polymerized for 15 min at room temperature followed by 30 min in the incubator (37°C, 5% CO_2_). Finally, the chips were supplied with cell culture medium via the side channels and incubated at 37°C with 5% CO_2_. Tumor spheroid growth from single cells proceeded for 7 days before nanoparticle incubation and PTT exposure.

### Nanoparticles synthesis and characterization

Iron oxide (Fe_3_O_4_) NPs coated with PO-PEG-NH_2_, 9nm core diameter, 30nm hydrodynamic diameter were produced by a non-aqueous sol-gel procedure as detailed in (47). Pegylation was modified by replacing POPEGCOOH by POPEGNH2 (specific polymers, catalog #SP-1P-14-001).

The core size of the nanoparticles (NPs) was determined from TEM images taken using a FEI CM10 electron microscope. Samples were prepared by dropping the colloidal solution onto a holey carbon-coated Cu grid. Hydrodynamic size and zeta potential were analyzed with a Nano ZS (red badge) ZEN3600 Zetasizer at neutral pH and 37°C. Infrared spectroscopy using KBr pellets was employed to qualitatively assess the efficiency of PO-PEG-NH_2_ coupling on iron oxide NPs. The FTIR spectra were recorded with a Bruker FTIR (Tensor 27) and are reported in terms of absorption frequency (cm^−1^). The 808 nm epsilon was determined from UV−visible spectra of aqueous NPs solution varying the iron concentration from 0.25 to 5 mM and was recorded using a spectrophotometer (V-630 UV−vis Spectrophotometer, Jasco). The crystalline composition of the iron oxide was determined using Mössbauer spectroscopy at 300K with the aid of a ^57^Co/Rh gamma-ray source in a transmission scheme with a triangular velocity form. The hyperfine structure was modelled by means of a least-squares fitting procedure involving Zeeman sextets composed of Lorentzian lines using MOSFIT (made home software). To describe the broadening of lines, several magnetic subcomponents (hyperfine magnetic field distribution) have been considered. The isomer shift (IS) values were referred to that of α-Fe at 300 K. Magnetic properties were assessed using a Vibrating Sample Magnetometer (VSM, Quantum Design, Versalab), with magnetization curves measured at 300K over a field range of –30,000 to 30,000 Oe.

For biological experiments, NPs were diluted in RPMI 1640 medium (Gibco 32404-014) supplemented with 1% of penicillin-streptomycin (P/S, Gibco 15140-122), and 10% fetal bovine serum (FBS, Gibco 10270-106) to prepare desired NP concentrations. Final concentrations of 8, 16 and 32mM were used. The NP solutions were then introduced into the chips via side chambers and left for 24h at 37°C with 5% CO_2_ to diffuse into the central chamber. The remaining NP solution was flushed with RPMI medium from the side chambers before laser exposure.

### Magnetite nanoparticle photothermal therapy (MagNP-PTT) setup

To implement nanoparticle-mediated photothermia heating in a microfluidic chip, the setup was developed. For this, the chip was first placed into a 3D-printed holder supplied with circulating warm water to bring the chip up to the initial temperature of 36.5-37.5°C before starting the laser exposure. An 808nm laser (Laser Components S.A.S France) is positioned in front of the chip at 6.1cm distance. This distance was identified as optimal to obtain a higher degree of homogeneity of laser surface power distribution that follows a Gaussian distribution profile while preserving high laser power range. In this study, the chips were exposed to the laser for 20min at 1 W/cm^2^ (3.12 A) to 2 W/cm^2^ (4.89 A). Temperature increase was recorded at 1.0Hz with an infrared thermal camera (FLIR A6751) and processed with FLIR ResearchIR Max software.

### Cell death rate evaluation

Cell survival rate was evaluated through confocal microscopy imaging, live/dead assay. LIVE/DEAD^TM^ Cell Imaging Kit (Invitrogen^TM^, R37601) solution prepared according to manufacturer’s instructions and incubated for 30min at 37°C with 5% CO_2_. Confocal microscopy (Leica DMi8) was performed with x25 magnification lens in water-immersion mode. The resulting images were obtained as 3D images (z-stacks, step size 2.0µm) with two channels: Calcein Green & Texas Red, identifying live and dead signal respectively. To analyse the images quantitatively, an algorithm was developed with an interactive machine learning tool, Ilastik 1.4.0rc6, to transform the images into probability maps of signal vs background. These images were further processed in ImageJ software by applying the threshold (P(signal) > 0.5) to convert the images into binary information and calculate the total area of signal in 3D. This was done for each 3D image, for live and dead signals separately. The ratio of red to green signals was then calculated as an estimate for cell death rate: 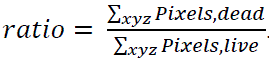.

### Collagen matrix visualization

To assess collagen fibrils upon NP-PTT exposure, collagen was stained with collagen hybridizing peptide, 5-FAM conjugate (F-CHP, 3Helix Inc., FLU200 / FLU60). For this, 5µM solution of F-CHP was prepared in PBS followed by heating at 80°C for 5 min in a water bath and quenching on ice for 60-90sec. The quenched solution was then introduced into the chips immediately and incubated overnight at 4°C protected from light. Next day, the chips were washed with 4 rounds of PBS of 30min each before imaging. Stained collagen fibrils were then imaged for fluorescence with an inverted confocal microscope (Leica DMi8) with x40 magnification lens in oil-immersion mode.

Alternatively, collagen fibers were visualized with second harmonic generation (SHG) microscopy using the inverted two-photon laser-scanning confocal microscope (Leica SP8) with a femtosecond laser (Chameleon Vision II, Coherent Inc.), Exc/Em 940/490 nm.

### Transmission Electron Microscopy

Self-formed spheroids in collagen on chips were analyzed by TEM to observe spheroids’ integrity and nanoparticle localization within the cells and the matrix as well as the impact of NP-PTT on spheroids. For this, spheroids were imaged either after extraction from collagen (via collagenase A digestion (1 mg/mL in DMEM+10%FBS+1%P/S, Roche Diagnostics GmbH)) or directly within the preserved collagen matrix. Spheroids (with or without the matrix) were washed with 0.1M cacodylate buffer (Sigma-Aldrich) and fixed with glutaraldehyde 5% followed by 2x washes with cacodylate buffer. Samples were further dehydrated, embedded in Epon, ultrasectioned for 70nm slices and placed onto copper grids. Finally, imaging was performed with a Hitachi HT7700 electron microscope operated at 80 kV (Elexience, France) with the help of MIMA2 MET – GABI, INRA, Agroparistech, 78352 Jouy-en-Josas, France.

### Prussian Blue (Perls’) staining

Intracellular ferric ions were visualized with the Prussian blue chemical reaction. Tumor spheroids in collagen on chip were initially fixed with 4% paraformaldehyde (PFA) for 30 minutes at room temperature. Iron staining was performed by treating the cells with potassium hexacyanoferrate (II) trihydrate (Sigma) under acidic conditions for 15 minutes. The nuclei were subsequently stained with Nuclear Fast Red solution for 15 minutes.

### Relative Quantification of Gene Expression by Real-Time PCR

Tumor cells were collected from either control (tumor spheroids grown on chip for 9 days), control+NPs (tumor spheroids grown on chip for 9 days with 24h NP incubation from D7 to D8) or post-NP-PTT at 40-44oC (tumor spheroids grown on chip for 9 days with 24h NP incubation from D7 to D8, laser exposure on D8 and 24h rest time). At least 6 independent samples were analyzed for each of the control conditions and at least 10 for post-NP-PTT condition per experiment, for each independent data point. Three independent experiments were performed. The total RNA was isolated using the NucleoSpin RNA mini kit (Machery-Nagel, Thermo Fisher Scientific, #740955.50) according to manufacturer’s instructions. Reverse transcription was carried out using the High Capacity cDNA Reverse Transcription kit (Thermo Fisher Scientific, #4368814) with random primers, following the manufacturer’s protocol. Both the extracted RNA and resulting cDNA were assessed for concentrations and purities using the Nanodrop device (Ozyme). Real-time quantitative PCR was performed using Power SYBRGreen qPCR Master Mix (Thermo Fisher Scientific, #A25742) on the QuantStudio 3 instrument (Applied Biosystems). The expression of Ribosomal protein P0 (RPLP0) expression was used as a reference gene to normalize the Ct values of the target genes. The normalized Ct values were then compared to those of the controls, and gene expression fold changes were calculated using the comparative Ct method: 2^−(ΔCtpostTreatment− ΔCtcontrols)^. Primer sequences are provided in Table S1.

### Immunofluorescence staining

The conditions stained include tumor spheroids in collagen on chip of: control (tumor spheroids grown on chip for 9 days), control+NPs (tumor spheroids grown on chip for 9 days with 24h NP incubation from D7 to D8) and post-NP-PTT at 40-44oC (tumor spheroids grown on chip for 9 days with 24h NP incubation from D7 to D8, laser exposure on D8 and 24h rest time). Subsequently, all samples were fixed on D9 with paraformaldehyde (PFA, 4% in PBS) for 45min at RT and washed thoroughly with PBS. They were then permeabilized with Triton (Triton X−100, 0.1% in PBS) for 20 min at RT and blocked with a blocking solution containing 4% bovine serum albumin (BSA) and 0.1% Triton X-100 in PBS for 3 hours at ambient temperature. The tumor spheroids on chip were then incubated with mouse anti-E-cadherin (1:200, Invitrogen SHE78-7) and guinea pig anti-Vimentin (1:400, ProGen GP53) primary antibodies in blocking solution overnight at room temperature. The following day, the samples were washed three times with PBS and incubated overnight at room temperature with Alexa Fluor 488-conjugated goat anti-mouse IgG (H+L) (1:400, ThermoFisher A11029) and Alexa Fluor 647-conjugated donkey anti-guinea pig IgG (H+L) (1:100, ThermoFisher A21450) secondary antibodies in blocking solution also containing 300 nM DAPI (D3571, Invitrogen) at 1:500 dilution. Finally, the chips were washed three times with PBS before imaging. Imaging was conducted in z-stack sections using a Leica DMi8 inverted confocal microscope (Leica Microsystems).

### Statistical analysis

All data distributions were first tested for variances homogeneity, and normality (D’Agostino-Pearson test, α=0.05). Accordingly, the Kruskal-Wallis nonparametric test was performed with multiple comparisons (Dunn’s correction, α=0.05). For two-factor analysis, Mann-Whitney two-tailed pairwise comparison was performed (α=0.05). All statistical significances are represented as *p < 0.05, **p < 0.01 and ***p < 0.001. All graphical representations and statistical analysis were performed with GraphPad Prism 9 software. The number of independent experiments was systematically greater than 3.

