## Supplementary information for "Magnetite Nanoparticle Photothermal Therapy in a Pancreatic Tumor-on-Chip: A Dual-Action Approach Targeting Cancer Cells and their Microenvironment"

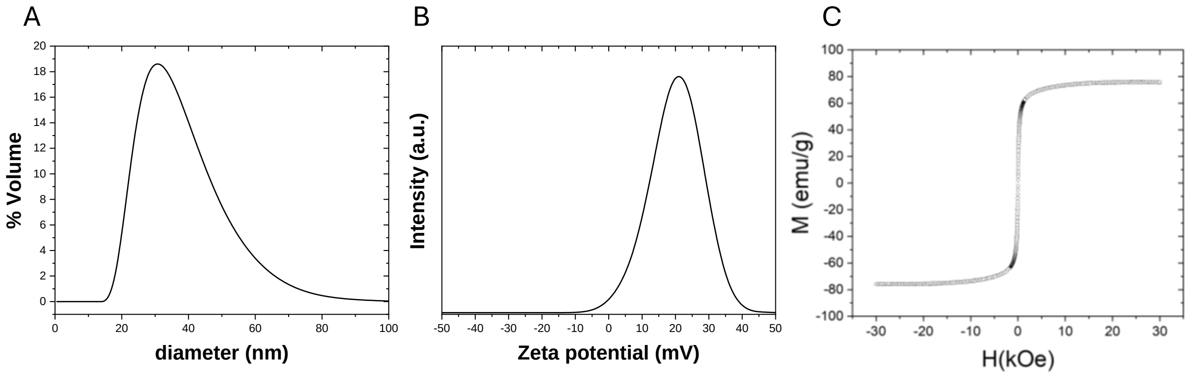


**Figure S1 :** $Fe_3O_4$@PO-PEG-$NH_3^+$ hydrodynamic diameter (A) and zeta potential measured at 37°C (B). (C) Magnetization curves at 300K.

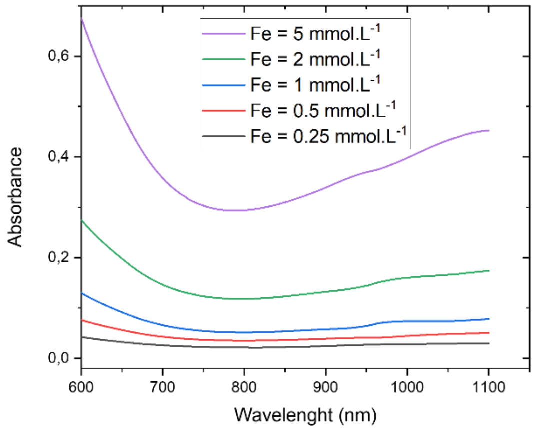


**Figure S2 :** Vis-NIR absorbance spectra of $Fe_3O_4$ NPs from 0.25 to 5 mM iron concentration.

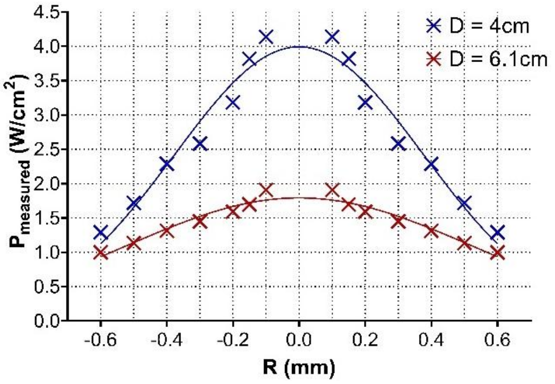


**Figure S3:** Laser surface power distribution at $P_{laser}$=2W at 6,1cm vs 4cm distance with Gaussian nonlinear fit through least squares regression.

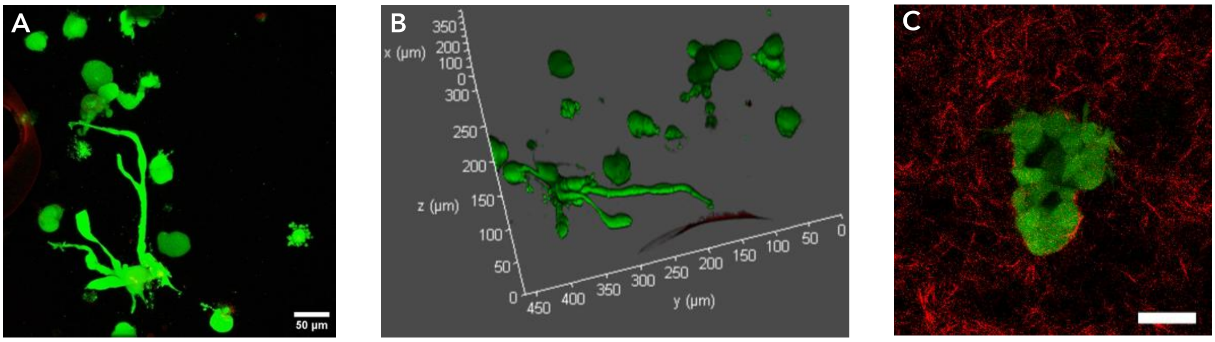


**Figure S4: 3D PDAC model in collagen type I on chip with characteristic protrusions.** Confocal microscopy images of Live/Dead assay ((dead = red; live = green)) of (**A**) z-projection of the 3D stack and (**B**) 3D representation on D8 of cancer cell-formed spheroids developed protrusions. (**C**) Second harmonic generation (SHG) microscopy image of a tumor spheroid (green) with the surrounding collagen type I matrix (red). Scale bar = 50µm.

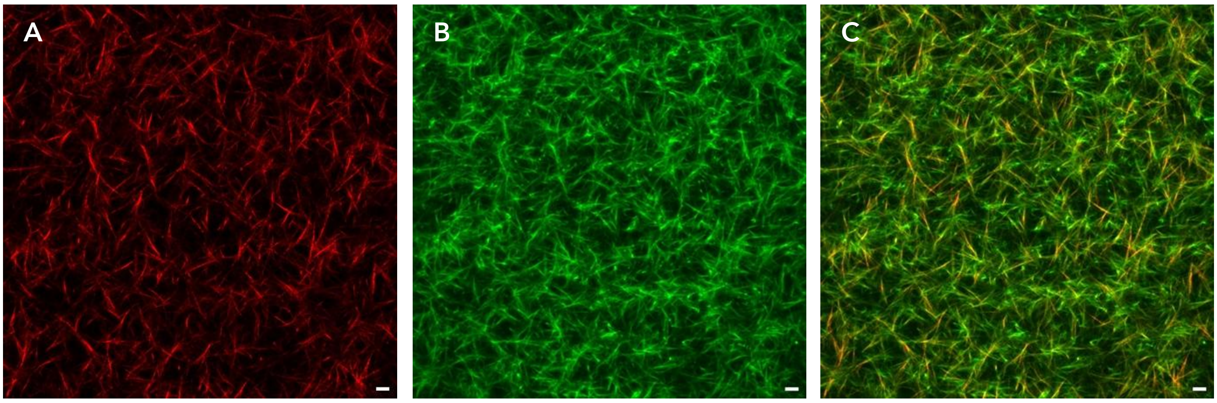


**Figure S5: Comparison of two-photon second harmonic generation (SHG) microscopy imaging with collagen hybridizing peptide imaged with confocal microscopy for collagen fiber visualization.** Collagen type I on chip at 6mg/mL is visualized with A. SHG signal. B. CHP signal. C. Overlap. Scale bar = 10µm.

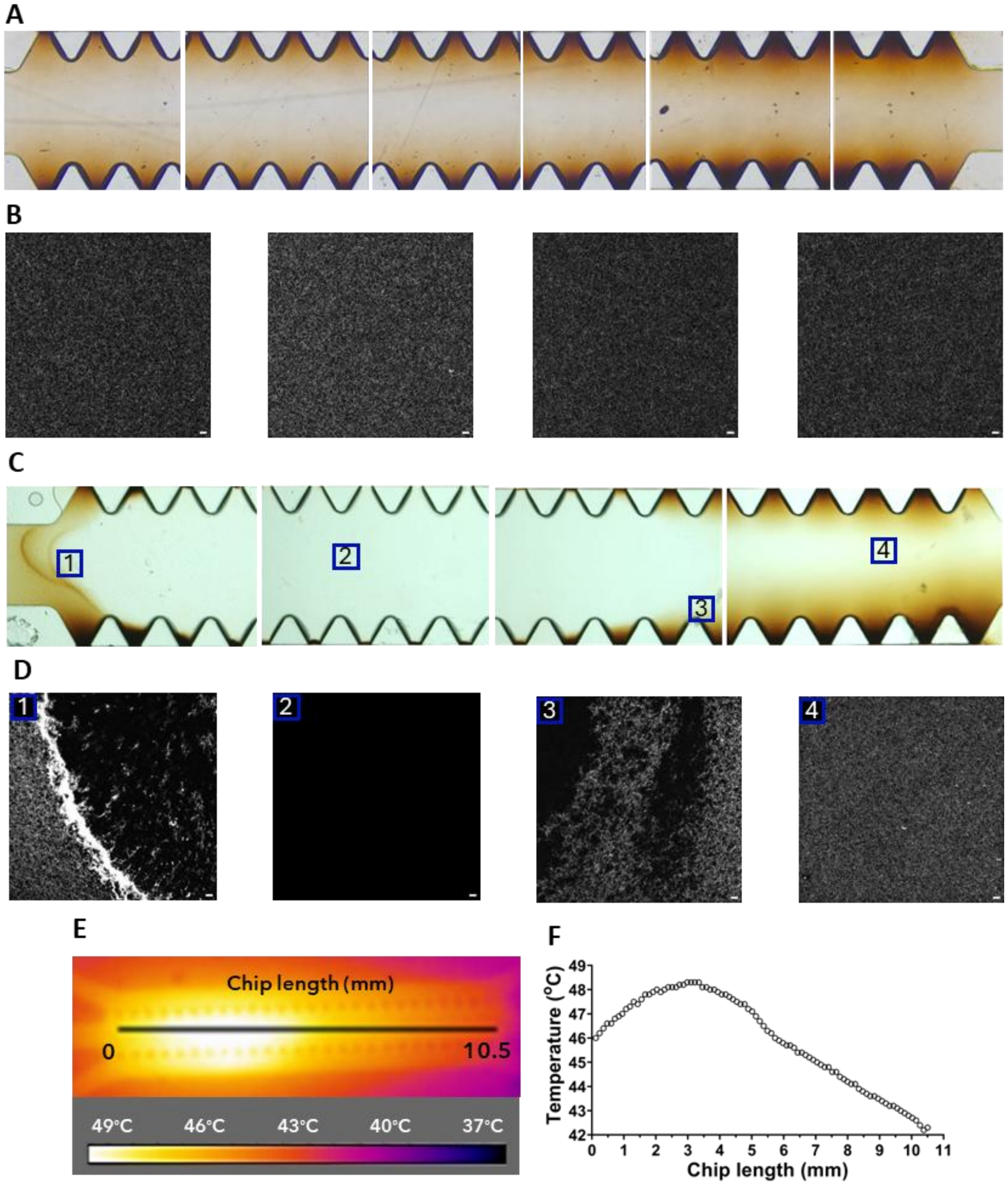


**Figure S6: Nanoparticle-mediated photothermia matrix impact characterization. A**. Bright field images of the control chip with NPs at 32mM diffused in collagen. **B**. Collagen hybridizing peptide staining imaging of different areas in the control chip with 32mM NPs with confocal microscopy. **C**. Bright field images of the chip (16mM NPs) with partially denaturated collagen. **D**. Collagen hybridizing peptide staining imaging of variously denaturated areas in the exposed chip with 16mM NPs, confocal microscopy. **E**. Corresponding thermal camera heating recording within the chip with representative temperature gradient along the length of the chip. **F**. Temperature variation as a function of the chip length

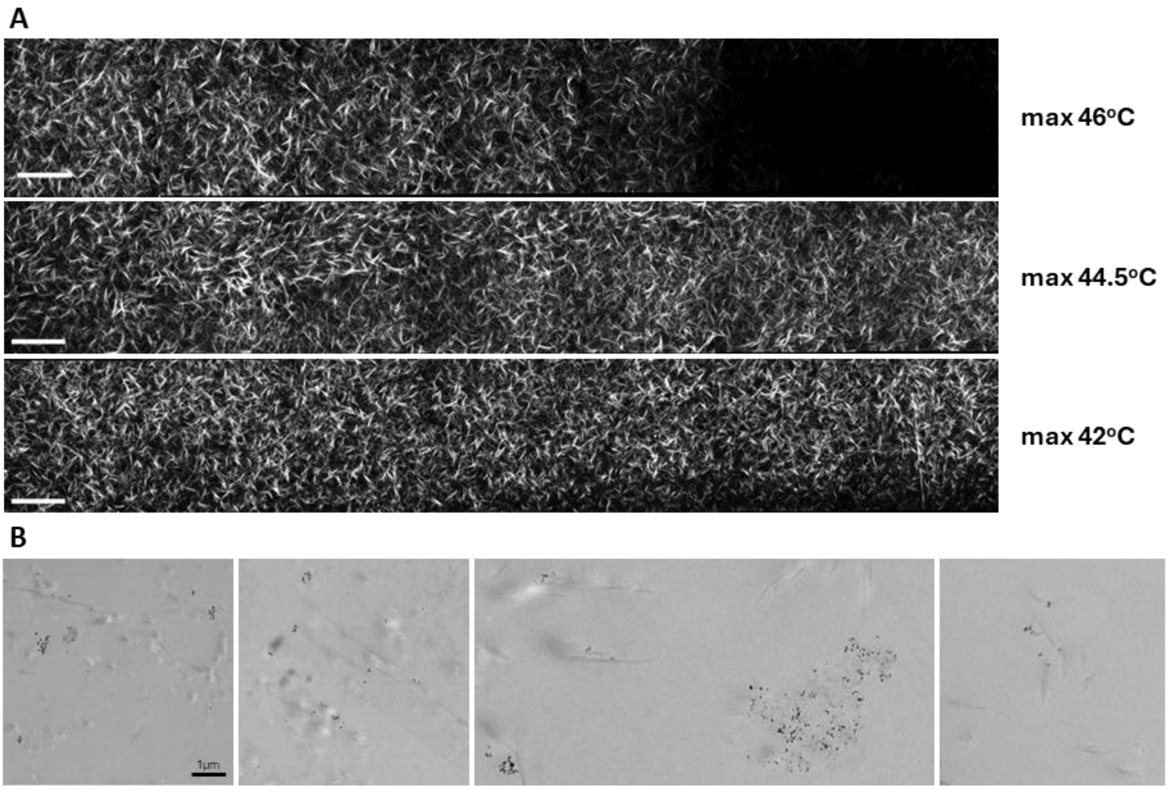


**Figure S7: Nanoparticle-mediated photothermia impact on collagen fiber structure. A.** SHG microscopy images of collagen denaturation following differential temperature exposure as a result of NP-PTT. **B.** TEM imaging of few remaining partially denaturated collagen fibers with adsorbed NPs (appear in dark contrast).

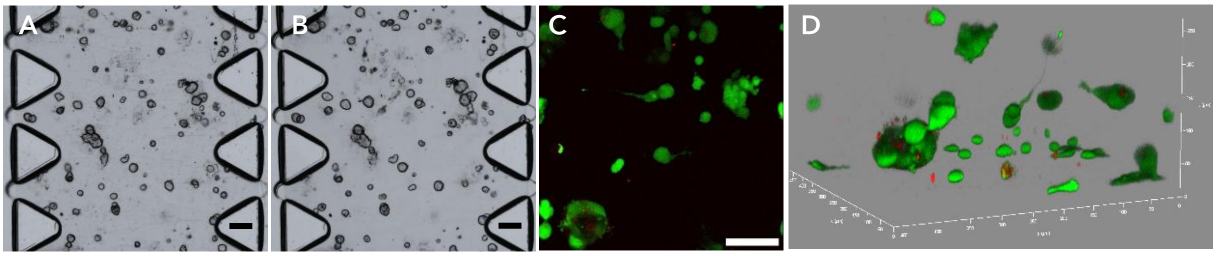


**Figure S8: 3D PDAC model in collagen type I on chip, control no NPs, 2W/cm$^2$ PTT.** A-B: Bright field microscopy images of control 2W/cm$^2$, on D7 before (A) and on D8 after (B) laser exposure. Scale bar = 100µm. C-D. Confocal microscopy images of Live/Dead assay ((dead = red; live = green)) of (C) z-projection of the 3D stack and (D) 3D representation on D8. Scale bar = 100µm.

**Table S1**

| **Gene name** | **Forward primer (5' to 3')** | **Reverse Primer (5' to 3')** |
|---|---|---|
| **BIRC5** | AGC CAA ATT CGT TGT CAT AC | GGT CTC CTC TGA CTT CAA CA |
| **MCL1** | CAG CGA CGG CGT AAC AAA C | ACA AAC CCA TCC CAG CCT CTT T |
| **Vimentin** | TAC AGG AAG CTG CTG GAA GG | ACC AGA GGG AGT GAA TCC AG |
| **ZEB** | TTC ACA GTG GAG AGA AGC CA | GCC TGG TGA TGC TGA AAG AG |
| **CDH2** | AGC CAA CCT TAA CTG AGG AGT | GGC AAG TTG ATT GGA GGG ATG |
| **FBN1** | GCT CCC AAA CCC TGC AAT TT | GGC AGT TGT GTT GCT TGG TTG |
| **FAK** | ATC CCA CAC ATC TTG CTG ACT T | GCA TTC CTT TTC TGT CCT TGT C |
| **CDH1** | ATT GCT CAC ATT TCC CAA CTC C | CTC TGT CAC CTT CAG CCA TCC T |
| **α-catenin** | GCG AAG GAG AGC CAG TTT CT | ATG TTG CCT CGC TTC ACA GA |
